# Benchmarking of analytical combinations for COVID-19 outcome prediction using single-cell RNA sequencing data

**DOI:** 10.1101/2023.01.18.524481

**Authors:** Yue Cao, Shila Ghazanfar, Pengyi Yang, Jean Yang

## Abstract

The advances of single-cell transcriptomic technologies have led to increasing use of single-cell RNA sequencing (scRNA-seq) data in large-scale patient cohort studies. The resulting high-dimensional data can be summarised and incorporated into patient outcome prediction models in several ways, however, there is a pressing need to understand the impact of analytical decisions on such model quality. In this study, we evaluate the impact of analytical choices on model choices, ensemble learning strategies and integration approaches on patient outcome prediction using five scRNA-seq COVID-19 datasets. First, we examine the difference in performance between using each single-view feature space versus multi-view feature space. Next, we survey multiple learning platforms from classical machine learning to modern deep learning methods. Lastly, we compare different integration approaches when combining datasets is necessary. Through benchmarking such analytical combinations, our study highlights the power of ensemble learning, consistency among different learning methods and robustness to dataset normalisation when using multiple datasets as the model input.

**Summary key points:**

- This work assesses and compares the performance of three categories of workflow consisting of 350 analytical combinations for outcome prediction using multi-sample, multi-conditions single-cell studies.
- We observed that using ensemble of feature types performs better than using individual feature type
- We found that in the current data, all learning approaches including deep learning exhibit similar predictive performance. When combining multiple datasets as the input, our study found that integrating multiple datasets at the cell level performs similarly to simply concatenating the patient representation without modification.

## Introduction

Single-cell RNA-sequencing (scRNA-seq) is a powerful tool that measures the transcriptomes of individual cells. As the technology advances, a typical dataset size has grown from a few thousand cells in 2014 to hundreds of thousands of cells in 2022 [1]. We are now in the era where the technology enables us to collect large pools of cells from multiple patients across multiple conditions. The current single-cell literature has mostly focused on analysing gene and cell level changes [2,3], for example dissecting the transcriptional heterogeneity in the population of single cells and identifying genes that mark the cell types [4]. Recently, there is an increasing number of studies designed at an individual level, such as between normal and patients (i.e. case vs control). These studies create the opportunity to examine disease mechanisms from multiple aspects, such as cell-type-specific changes in gene expression, pathway regulation [5,6] and cell-cell interaction [7] in order to gain a deeper understanding of disease mechanism. The analysis of such data requires the development of methods that can extract information from multi-sample multi-condition disease study designed at an individual level rather than at a cell level [8].

To date, the majority of the questions at individual level have focused on the identification of differentially expressed genes between cell types and states [9], and differential abundance of cells between states [10,11] of individuals. A natural next question is to develop models that explore at a higher level how the outcome associated with each individual can be predicted in multi-sample multi-condition scRNA-seq dataset. As the number of individuals increases, there will be a demand to develop models that accurately predict the outcome of each patient in such data. To meet this demand with an interpretable focus, it will be necessary to first extract informative features from complex single-cell data structures that represent each individual and then understand which approaches are most effective for utilising the summary statistics for downstream analysis.

To date, a large repertoire of approaches has been developed to model for the prediction task, which prompts the question: “What are the optimal approaches?”. Since the past decade, modern deep learning has gained tremendous success compared to classical machine learning in analysing complex data. However, it is worth noting that deep learning models often involve millions of parameters [13] and require larger amounts of data and computational resources to train compared to classical machine learning.This raises the question of whether it is necessary to use deep learning models. In parallel, when extracting information from data, we may obtain multiple pieces of information. The fusion of multiple information, or ensemble learning [14], is a common technique to improve the performance of prediction model. There are various ways to fuse the information [15], including at the input feature level, at the model level and at the predicted outcome level. The question here is whether ensemble of features improves performance and which ensemble strategy is the most optimal. The increasing availability of single-cell datasets has led to the availability of multiple datasets focusing on the same interest, such as a particular disease. This strategy naturally lends itself to combining multiple datasets and enabling investigation that may not be possible with a single dataset. An important question to address is how to optimally integrate these datasets to achieve the best performance.

In this study, we examine the question of the optimal approaches as mentioned above using uniquely collected COVID-19 scRNA-seq datasets. To generate derived statistics for each patient sample, we utilise the recently developed scFeatures [8] method that constructs a multi-view representation across various feature types. We implement and compare different learning models from classical machine learning to modern deep learning models. We compare the performance of individual feature types as well as the ensemble of feature types by implementing a number of common ensemble strategies [14,16]. Additionally, using multiple COVID-19 datasets, we investigate the optimal data integration approach that maximises prediction outcomes. Overall, through a comparison framework, we assess the combined impact of these key data analytical components (i.e., model choices, ensemble learning strategies and integration approaches) on COVID-19 outcome prediction.

## Material and Methods

### Design of a benchmarking study

Any benchmarking or comparison study typically involves three key elements. First, a collection of datasets is needed to evaluate the performance of methods without bias. Second, well-designed evaluation strategies are needed to compare methods or workflows. Thirdly, evaluation metrics from multiple aspects are needed to quantify the performance. These elements are described in more detail in the following sections.

### Evaluation datasets collection

As the world has been heavily impacted by COVID-19 for over three years, the global effort in understanding this pandemic has made COVID-19 patient data perhaps one of the largest collections of multi-sample multi-condition single-cell datasets to date. Therefore, to examine the data analytics strategy for cohort analysis, we selected five large publicly available COVID-19 datasets containing individuals with mild and severe disease progression (Figure 1a). This collection includes a total of 2,215,517 cells from 223 mild and 245 severe patients. All datasets are composed of peripheral blood mononuclear cells (PBMC) or whole blood samples. Additional details are provided in Table 1.

**Table 1.** Collection of COVID-19 PBMC datasets sequenced using scRNA-seq providing a total number of 2 million cells and 471 individuals.

| Dataset name | Reference | Accession ID | Number of mild individuals | Number of severe individuals | Number of mild and severe individuals | Number of cells in mild and severe individuals |
| --- | --- | --- | --- | --- | --- | --- |
| Combat | [20] | EGAS00001005493 | 30 | 61 | 91 | 524,557 |
| Ren | [21] | GSE158055 | 68 | 85 | 153 | 872,663 |
| Schultes-schrepping | [22] | EGAS00001004571 | 44 | 51 | 95 | 212,023 |
| Stephenson | [23] | E-MTAB-10026 | 58 | 32 | 90 | 493,685 |
| Wilk | [24] | GSE174072 | 23 | 19 | 42 | 112, 589 |
| Total |  |  | 223 | 245 | 471 | 2,215,517 |

**Figure 1.**
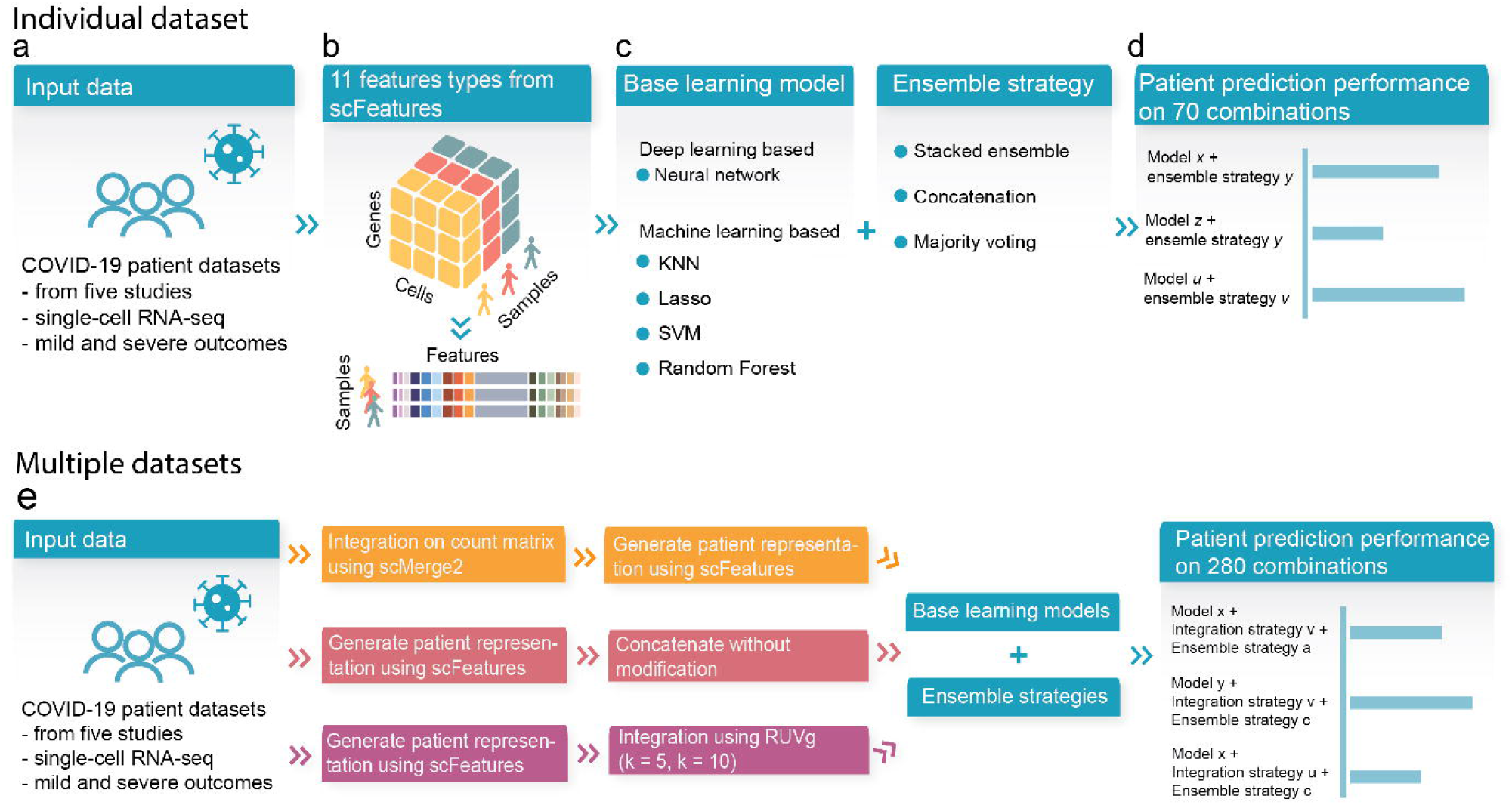
Schematic of the benchmark workflow. **a** Five COVID-19 scRNA-seq datasets containing mild and severe outcome patients were used in this benchmark study. **b** We used scFeatures to generate 11 types of molecular representations of each individuals (i.e., the patients). **c** We implemented five models containing both deep learning and machine learning, as well as three ensemble strategies. **d** The analytical strategies resulted in a total of 70 combinations for evaluating patient outcome prediction in each individual dataset. **e** We also evaluated the performance of analytical strategies on the combined dataset. To combine the dataset, we implemented three integration strategies. We used the same base learning models and ensemble strategies as shown in **c**. This resulted in another 280 combinations.

### Evaluation strategies for analytical combinations

The comparison study aims to examine the impact of various analytical strategies on individual level outcome prediction. To accomplish this, we utilised our recently developed feature engineering tool, scFeatures, to generate multi-view molecular representation of each individual that served as input for downstream analytical models (Figure 1b).

The evaluation is composed of three key components (Table 2): (1) comparing the performance of multiple learning models using the generated features as input, (2) comparing single-view and multi-view features through ensemble strategies, and (3) comparing integration strategies when using multiple datasets as the input. In component 1, we surveyed and implemented multiple learning models from classical machine learning to modern deep learning methods (Figure 1c). In component 2, we examined the difference in performance between using single-view feature space versus multi-view feature space via implementing multiple ensemble strategies (Figure 1d). In component 3, given that we collected multiple COVID-19 datasets, we compared the performance of analytical choices on the combined dataset.

**Table 2.** Summary of the analytical choices implemented in this comparison study.

| **Analytical component** | **Analytical choice** |
| --- | --- |
| **Base learning model** | (1) KNN,<br>(2) Lasso,<br>(3) Random Forest,<br>(4) SVM,<br>(5) Neural network |
| **Ensemble strategy** | (1) Concatenation,<br>(2) Majority voting,<br>(3) Stacking |
| **Level of integration** | (A) Cell level integration,<br>(B) Individual level integration with no normalisation,<br>(C1) Individual level integration with RUVg adjustment (using $K = 5$),<br>(C2) Individual level integration with RUVg adjustment (using $K = 10$) |

On each of the individual datasets, we examined a total of 70 analytical combinations from 11 feature representations, five base models and three ensemble strategies (as detailed in Table 2). On the combined dataset, we applied the same 70 combinations to each of the four integration strategies, resulting in a further 280 combinations. Further details on each of the three components are given in the following subsections.

#### Feature generation

We used scFeatures to generate the molecular representation for each individual in each of the COVID-19 datasets. A total of 11 feature types from five feature categories were generated to reflect different views of the molecular property and were used for downstream analysis. In detail, the following feature representations were generated for each patient: (1) Proportion ratio, (2) Proportion raw, (3) Proportion logit, (4) Gene mean celltype, (5) Gene proportion celltype, (6) Pathway gsva, (7) Pathway mean, (8) Pathway proportion, (9) CCI, (10) Gene mean aggregate and (11) Gene proportion aggregated. Information regarding each of the feature types can be found in [8].

#### Base model selection

To examine the effect of different learning models on individual level outcome prediction, we examined a selection of approaches from classical machine learning methods to the more recent deep learning approach. In the rest of the paper, we used the word “machine learning” in its broadest definition to refer to both classical machine learning and deep learning methods.

For **classical machine learning approach**, we included a range of models including KNN, Lasso, Random Forest and SVM with linear kernel using the implementation in the R package Caret [17]. Each feature type was used individually as the input to compare the performance of each feature type. The severity (mild and severe) of the individuals’ conditions was used as the outcome variable. For Lasso which outputs the prediction in terms of probability instead of discrete outcome, we used 0.5 as the threshold.

For the representative **deep learning approach**, we implemented a neural network structure containing four fully connected layers. For each feature type, we used the same network structure but varied the number of nodes in the layers depending on the number of features in the feature type. In detail, the input layer had a number of nodes equal to the number of features in the respective feature type. The second layer and third contained different numbers of nodes depending on the feature types. We describe the detailed implementation below:

- All feature types in the category “cell type proportions” contained less than 100 features. For these feature types, we set both the first layer and second layer to 20 nodes.
- All feature types in the category “cell type specific pathway expressions”, “overall aggregated gene expressions” and “cell-cell communications” contained less than 1000 features. For these feature types, we set the first layer to 500 nodes and the second layer to 100 nodes to reduce the dimension.
- All feature types in “cell type specific gene expression” contained less than 10000 features. To reduce the dimensions for these feature types, we set the first layer to 1000 nodes and the second layer to 100 nodes.

The number of nodes in the output layer was the same for all feature types, with two nodes that output the probability of mild and severe conditions, respectively. The condition with higher probability was considered the predicted condition.

#### Ensemble strategies for multi-view features

We considered three types of ensemble strategies. scFeatures generates multiple feature types for a given patient, representing different and possibly complementary biological information (views). It is therefore of interest to examine the performance of ensemble learning by integrating multiple feature types (i.e., “multi-view”) compared to using each of the feature types individually as “single-view”. Here, we employed three types of ensemble strategies to obtain a “multi-view” prediction. We used the term “ensemble” in its broadest definition to refer to integrated learning at either feature or model level. Specifically, the implementation of these ensemble strategies is as follows:

1. **Early fusion** using **concatenation**, which involved concatenating features across all feature types as the input. The implementation was the same for both machine learning and deep learning models as this strategy operates on the feature level.
2. **Late fusion** using **stacked ensemble**. This involved training each base learner on a single-view of the feature space followed by training a meta-learner to best combine the individual base learners. The implementation was different for machine learning and deep learning models. For machine learning models, base learners were trained and evaluated on each of the individual feature types, resulting in 11 predictions for each patient. The predictions were then used as the input to build a logistic regression model, which serves as the meta-learner that combines the base learners to produce the final predicted outcome. For deep learning models, we implemented a network (Supplementary Figure 1) containing 11 subnetworks that took each of the 11 feature types as input. The subnetwork performed feature extraction for each of the feature types individually. We used the same network structure as the network described in the previous section that was used for extracting features from each feature type individually. The extracted features from each feature type were then concatenated, resulting in a vector of 860 features for each individual. This feature vector was then passed through a fully-connected layer containing 50 nodes, followed by the output layer containing two nodes to produce the final prediction.
3. **Score fusion** using **majority voting**. We first obtained the predicted outcome from each of the 11 feature types, resulting in 11 predictions of either mild or severe for each patient. Then the outcome with the most votes was considered to be the final predicted outcome for the patient. The implementation was the same for both machine learning and deep learning models.

#### Levels of integration strategy

We examined different levels of integration to explore the optimal choice for predicting patient states when multiple datasets need to be combined and used as a whole in building a prediction model. The approaches are described in the following:

- **Cell level integration -** this approach refers to **integration on count matrix**: We used scMerge2 (personal communication) to perform data integration on the scRNA-seq count matrix. We then generated the patient representation using scFeatures on the integrated count matrix and used this as input for learning model.
- **Individual level integration with no adjustment:** We simply concatenated the patient representation without any adjustment or normalization, and used this as input for learning model.
- **Integration on patient representations**: We used a well-known batch correction method RUVg [18] to correct for the batch effect in the patient representation. As *k*, the number of unwanted variations is a tunable parameter, we explored two settings of k = 5 (i.e., where the number of batches is equal to the number of datasets) and k = 10 (i.e., to introduce a stronger batch correction). The batch-corrected patient representation was used as input for learning model.

### Evaluation metric

#### Accuracy metric

To quantify the performance of the methods, we recorded the prediction accuracy of the severity outcome (Figure 1e). To capture the variability in model performance, all classical machine learning and deep learning models were trained and tested with repeated three folds cross-validation using 20 repeats. To control for the potential impact of “good” or “bad” training/testing set splits, where a “bad” split can result in extreme class imbalance in the modelling phase and affect model performance, we used the same training and testing splitting index across all machine learning and deep learning model to ensure a fair comparison. F1 score was used as the evaluation metric, as not all datasets are balanced.

#### Aggregation of accuracy metric

Given the number of results from all analytical combinations, we aggregated the results in order to better quantify and interpret the results. First, we took the median F1 score across the 20 repeated cross-validation. This was then followed by different aggregation strategies depending on whether the input used individual or combined datasets.

For the result section where we dealt with the five datasets individually, we further aggregated the median F1 score across datasets by taking the median. Then, we ranked the feature types across each model choice as well as the model choice across each feature type to derive the ranking of feature types and the ranking of model choice.

#### Computational resource metric

Apart from assessing the performance in terms of accuracy, we also assess the performance in terms of the computational resources. This was measured through running time and memory usage averaged over three repeats. All processes were executed using a research server with dual Intel(R) Xeon(R) Gold 6148 Processor with 40 cores, 768 GB of memory and two NVIDIA GeForce RTX 2080 Ti graphics cards.

Running time of each combination was measured using the Sys.time function built in R and the time.time function built in Python. Memory usage was quantified in terms of CPU memory for combinations involving machine learning models. For combinations involving deep learning models, the memory usage was quantified as the sum of CPU and GPU memory, as the deep learning models were executed on GPU.

## Results

### Certain ensemble strategies improve model performance

In this study, we examined the impact of using ensemble of feature sets for predictive modelling in large cohort single-cell data by using scFeatures to extract 11 feature types for each individual. We compared the performance using prediction accuracy on patient outcomes across five COVID-19 patient datasets (Table 1, Supplementary Figure 2). Across the five machine learning approaches, we observed that certain ensemble strategies performed better than models based on individual features. In particular, majority voting consistently achieved the best performance, outperforming the other two ensemble strategies, as well as all individual features (Figure 2). This was followed by concatenation, which also performed better than using any of the individual features. These results highlight the effectiveness of ensemble learning and also suggest that the feature types are diverse, such that different feature types make different errors such that combining them leads to improved model performance. Further examination of the top eight learning model and feature type combinations (Figure 3) revealed that seven of the eight combinations involves ensemble learning. Interestingly, the more complicated implementation of ensemble learning called stacked ensemble, in which a meta-learner is trained on the base learners trained from individual feature types, performed worse than using any of the individual feature types except for when deep learning was used.

**Figure 2.**
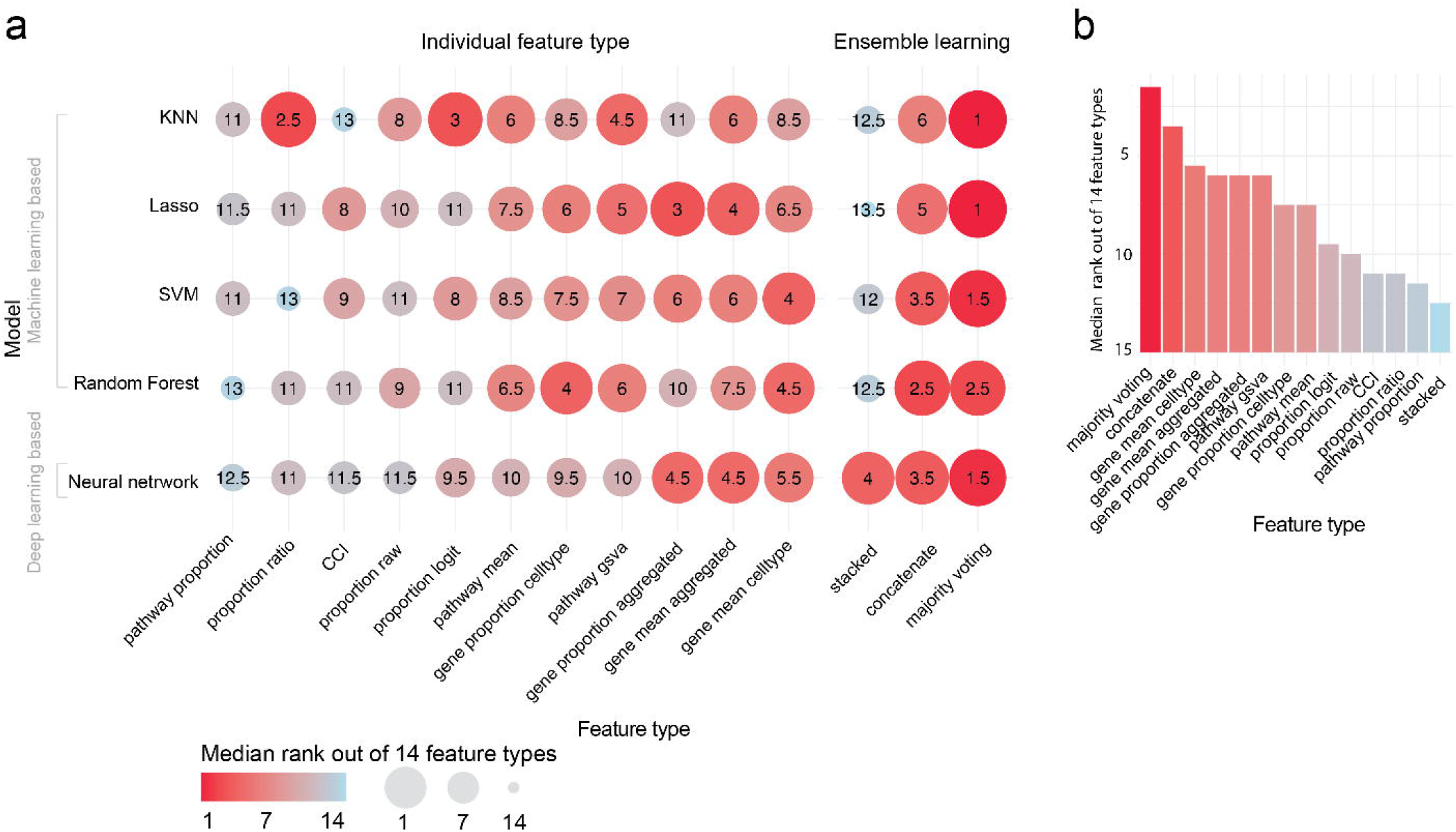
Performance of feature types for each model summarised across all datasets. The dotplot shows the relative rank of each feature type to each other for each model, with 1 being the best and 14 being the worst. Ranks are summarised across the five datasets using the median and therefore do not necessarily range from 1 to 14 within each model.

**Figure 3.**
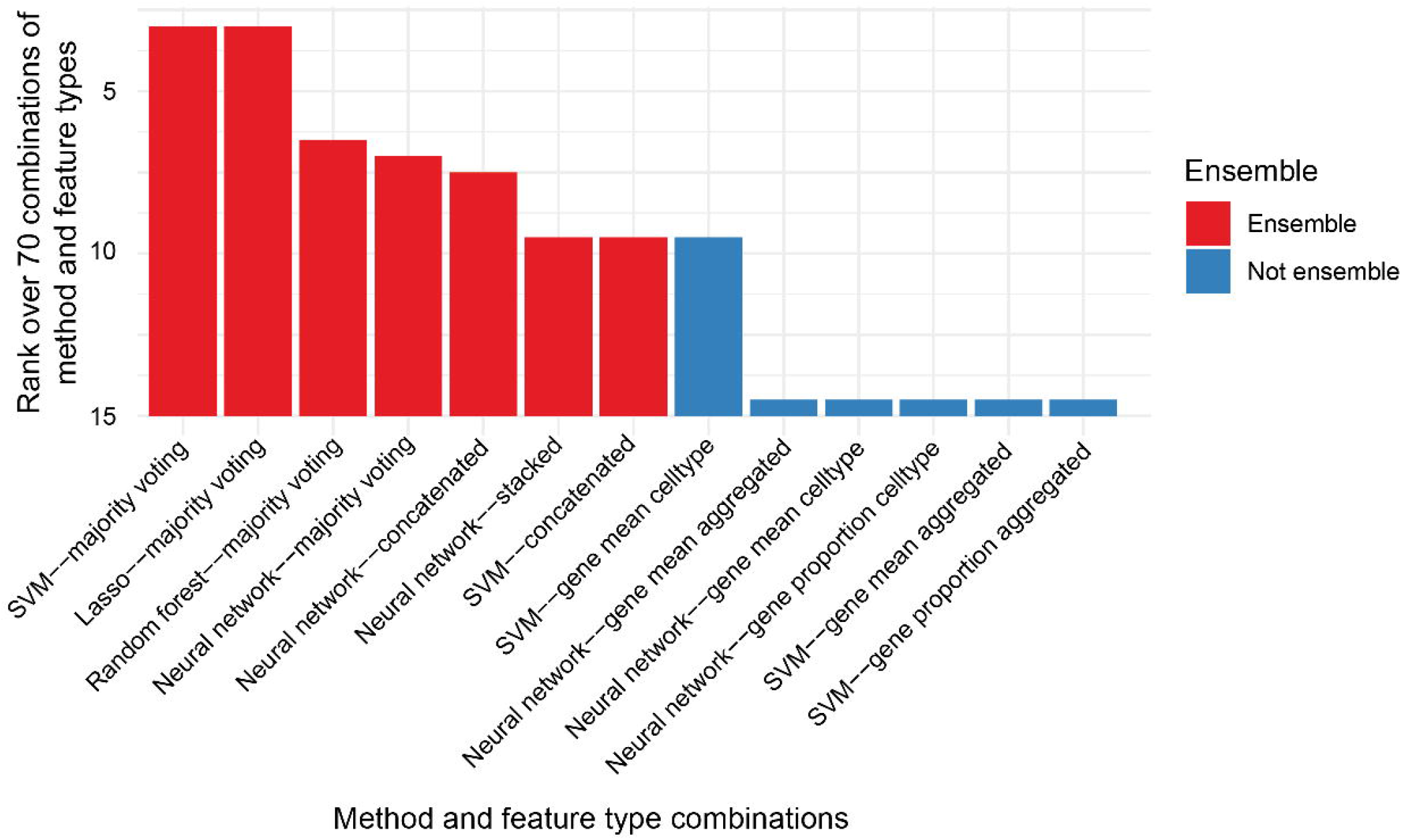
Top 13 combinations of model and feature type. Barplot shows the ranks of model and feature type. Given that the ranks are summarised across all five datasets using the median, the values do not necessarily range from 1 to 13.

We then took a closer examination at whether this observation is consistent irrespective of the learning model choice or dataset. We ranked the feature types on each of the five types of models and each of the five datasets. We observed that no individual feature type consistently ranked better or worse than others across all models and datasets (Figure 4). Almost all individual feature types had ranks that varied from 1 (the best rank) to 14 (the worst rank). This suggests that different feature types are useful for different models and different datasets, despite them all being COVID-19 datasets with mild and severe individuals. In contrast, majority voting achieved a rank of 1 across multiple models and multiple datasets, again illustrating the power of ensemble strategy.

**Figure 4.**
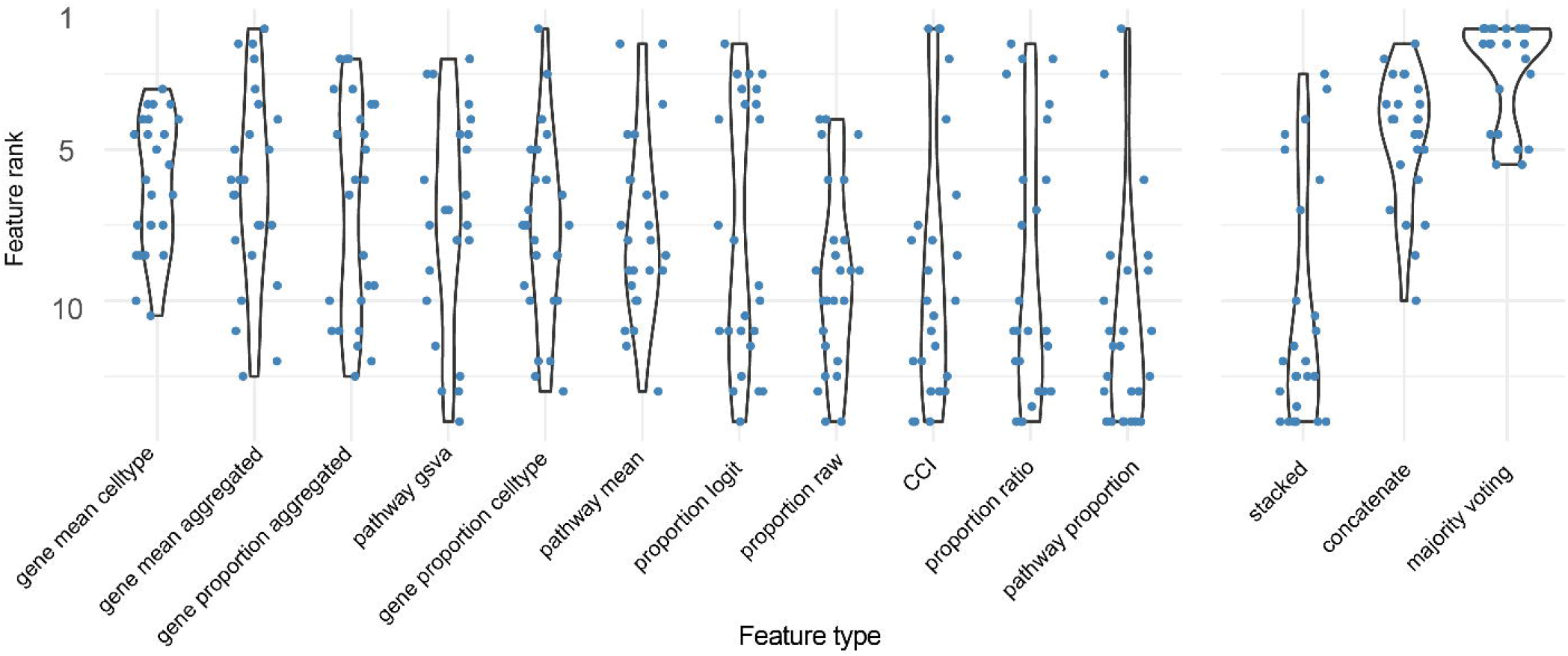
Performance of feature types for each model and each dataset. Violin plot shows the distribution of rank of each feature type for each model choice and each dataset choice. A total of 25 points are shown for each violin plot, as each feature type was evaluated on five models and five datasets.

### Deep Learning performs similarly to classical machine learning

Ranking the learning methods, we noted that there was no clear difference between deep learning and some of the machine learning models (Figure 5a). In particular, both neural network and random forest achieved a median rank of 1.75 out of the five learning methods across the 14 feature types and five datasets, followed closely by SVM with a median rank of 2.5 (Figure 5b). Only lasso and KNN were consistently ranked lower than other methods. Within neural network, random forest and SVM, we then examined the difference between the maximum and minimum F1 score achieved by the three top-performing methods and observed a small median difference of 0.02 (Supplementary Figure 3). These result all suggest that deep learning do not significantly outperform certain machine learning models in this context.

**Figure 5.**
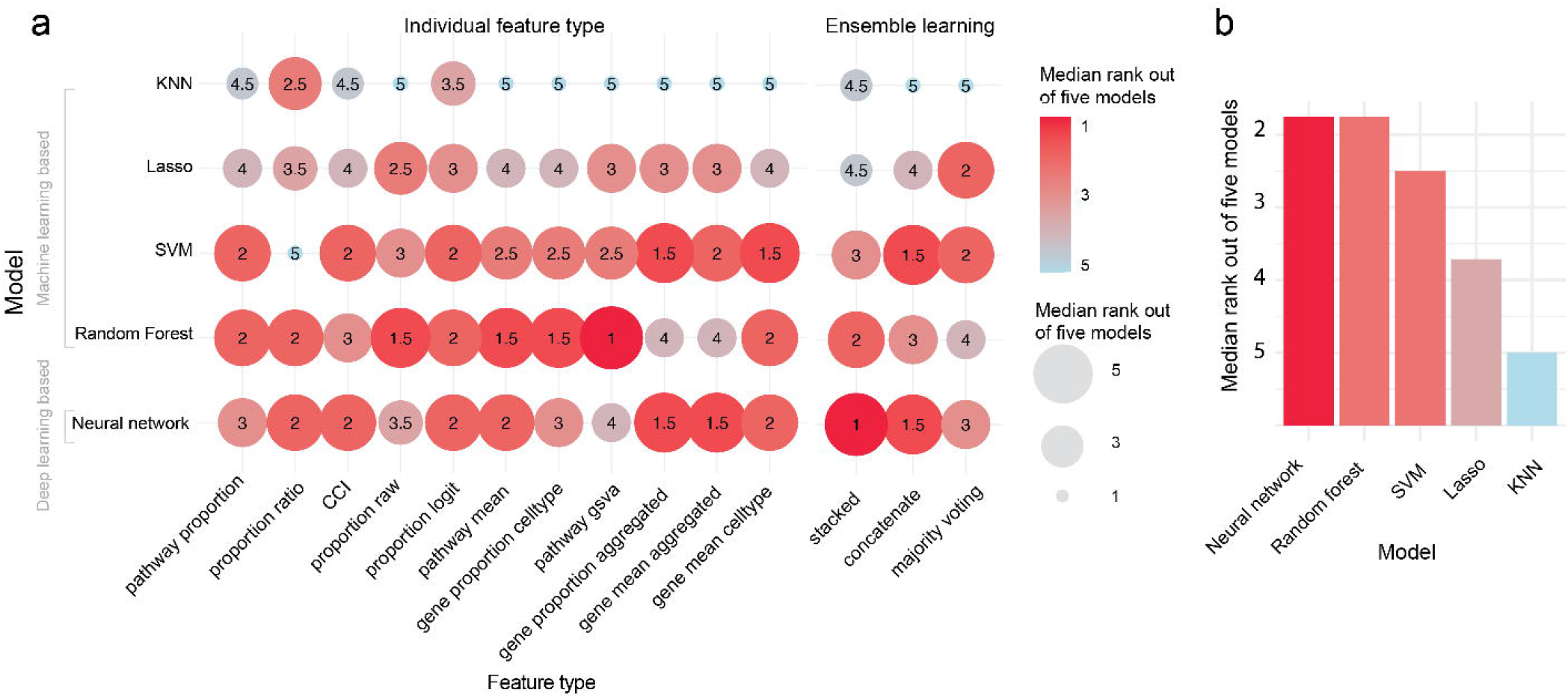
Performance of models. **a** shows the relative rank of each model to each other for each feature type with 1 being the best and 5 being the worst. Ranks are summarised across the five datasets using the median and therefore do not necessarily range from 1 to 5 within each feature type. **b** further summarise the ranks of each model across all feature types using the median.

We then compared the computational resource requirement to see whether the difference in performance came at a cost. Focusing on the feature type “majority voting”, we observed that both neural network and random forest took around 4 hours on the largest Ren et al. dataset with 153 patients (Supplementary Figure 45). On the other hand, while SVM was ranked after neural network and random forest, it was more computationally efficient, taking less than 1 hour on the Ren et al. dataset. Accounting for the significant difference in computational efficiency and the relatively small difference between model performance, one may consider SVM to be the optimal choice.

### Normalisation is not necessary when combining multiple datasets as the input

Using multiple datasets as input data raises a number of questions, such as whether to integrate the raw data or the derived features. Here, we explored three categories of analytical combinations. More specifically, different approaches to data integration, including integration on the count matrix, integration on the patient representation with and without normalization were explored. Our results are based on the examination of a total of 280 analytical combinations (4 integration types x 14 feature types [11 individual feature types with 3 ensemble feature types] x 5 model choices). Interestingly, there was only a slight difference between integration on the count matrix and concatenation without modification (Figure 6), which both achieved high F1 scores. On the other hand, integration on the patient representations achieved lower F1 scores, with the stronger the batch removal setting, the worse the F1 score. This observation is consistent across the choice of method and the type of feature used (Supplementary Figure 6 7).

**Figure 6.**
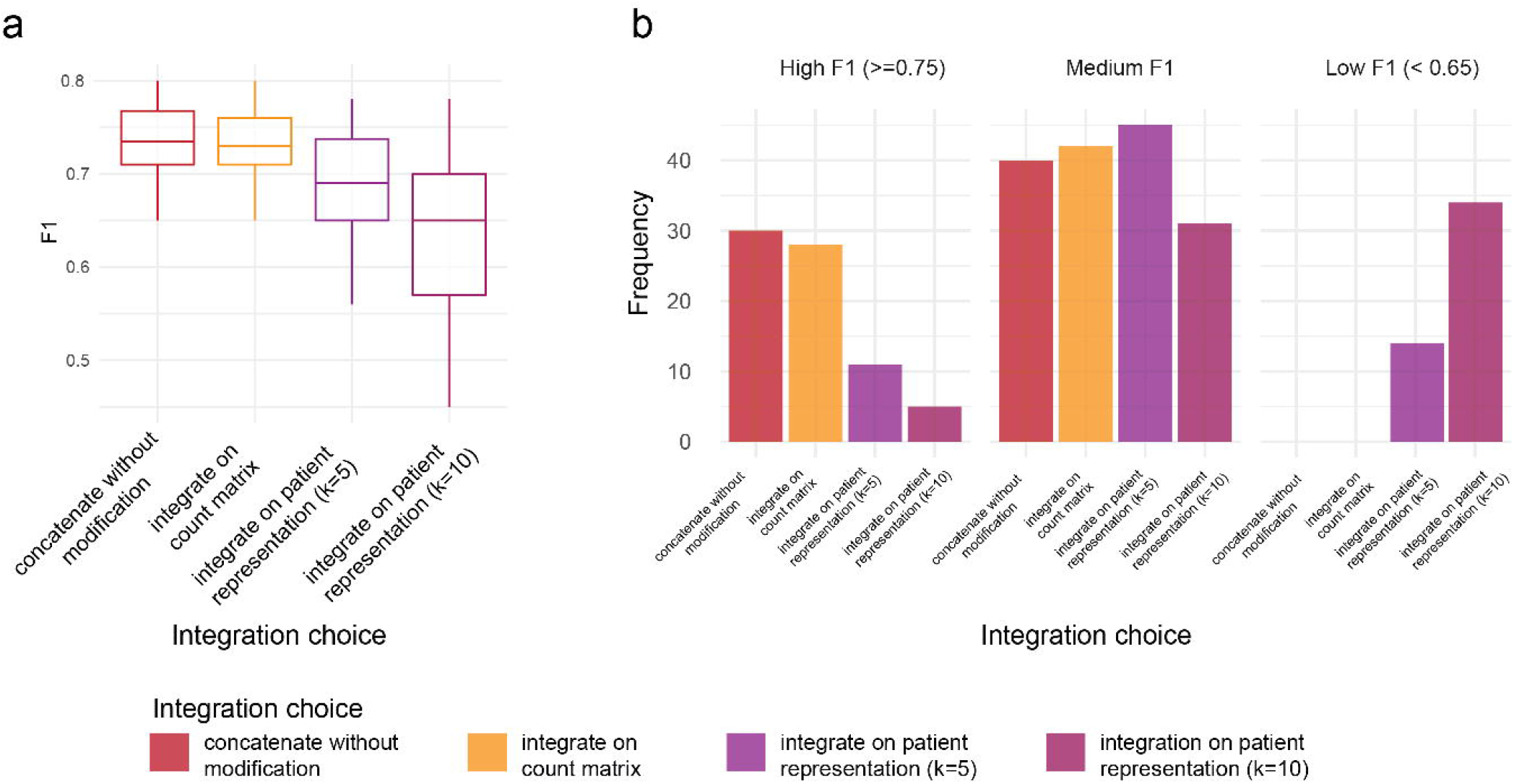
Performance of various approaches on combining multiple datasets for building prediction model. **a** shows the F1 of these 280 analytical combinations, with the x-axis indicating the type of integration choice used in the combinations. **b** further stratifies the F1 score based on high F1 (defined to be F1 >= 0.75), medium F1 score (defined to be 0.65 < F1 < 0.75) and low F1 score (defined to be F1 <=0.65) and examines the proportion of each integration choice in the set of combinations that fall in the stratification.

One of the key strengths of data integration is the ability to examine condition associated features for a subgroup of individuals. Due to the small number of individuals that typically fall into the subgroup of interest, this type of research is difficult to conduct using a single dataset. Here, we focused on a subgroup of patients in the 41-50 age group and investigated whether the identification of features is affected by different data integration strategy. First, we compared the rankings of the features obtained according to the feature importance score from the prediction model and found high consistency of the rankings between cell level integration and individual level integration without normalisation (Supplementary Figure 8a). In comparison, the consistency was much lower between cell level integration and individual level integration with normalisation. Clustering and dimension reduction on the features revealed that in both cases the clustering patterns and sources of variation of the patients were not driven by the dataset source (Supplementary Figure 8b,c). The lack of batch effect in the generated features suggests that the generated features may have self-adjusted in the feature extraction procedure, therefore explaining the minimal difference observed between the feature rankings and suggesting that there is no need for normalisation on cell level or on individual level.

## Discussion

In this comparison study we explored different analytical approaches for predicting the severity of COVID-19 using multi-sample multi-condition scRNA-seq data. We used scFeatures to generate various feature representations for COVID-19 patients and examined the performance of individual feature types and ensemble feature types in classifying COVID-19 severity. By evaluating using multiple datasets and multiple learning methods from classical machine learning to modern deep learning methods, this study demonstrated that all machine learning methods perform similarity, with SVM being a slightly better method when accounting for the computational efficiency. Through implementing different ensemble strategies to incorporate multiple feature types as input into machine learning models, we revealed certain ensemble strategies, in particular majority voting, consistently led to increased performance compared to the non-ensemble strategy of using individual feature types alone. Stacked ensemble for example, often did not achieve better performance compared to using individual feature types. Finally, we suggest that when combining datasets is required for a prediction model, prior data integration is not necessary and doesn’t necessarily improve prediction performance.

We observed that with the sets of COVID-19 datasets containing 42 to 153 patients, which is a realistic sample size in the current literature, the more complex approaches do not necessarily outperform simpler approaches. In particular, stacked ensemble can be considered the most complex implementation as it trains additional meta-learner on top of the base models. We observed that while the other two implementations (majority voting and concatenation) both performed better than individual features, stacked ensembled had worse performance compared to using the individual features. Furthermore, we observed minimal improvement when loading all single-cell data, a total of over 2 million cells for five datasets, as opposed to first generating patient-level features. With the extensive, and potentially prohibitive, computational resources required for such cell-level integration [19], such gain in model accuracy may not be worth the tradeoff in computational resources.

Recently, there have been a growing number of multi-sample multi-condition datasets. While this opens opportunities for patient level analysis such as case control study, it also opens new questions on what are the representative methods for each question, what are the appropriate quantitative evaluation metrics to assess each method, what are the recommended approaches for answering a given question with data of certain characteristics and what are guidelines for future methods development. In this study, we utilised five COVID-19 patient datasets to evaluate the choices of the method, ensemble strategy and integration strategy and obtained consistent trends. We envisage the current comparison framework will point valuable direction into an optimised analytical combinations for outcome prediction using single-cell data in future where cohort study with more than a few hundred or over a thousand patients are readily available.

## BACK MATTER

### Code and data availability

The data that support the findings of this study is publicly available and the accession ID is reported in Table 1. The scFeatures package used for extracting patient representation is freely available at our Github page: https://github.com/SydneyBioX/scFeatures.

## Acknowledgements

The authors thank their colleagues at The University of Sydney, Sydney Precision Data Science Centre, School of Mathematics and Statistics, Charles Perkins Center for intellectual engagement.

