## Supplementary Materials for "Benchmarking of analytical combinations for COVID-19 outcome prediction using single-cell RNA sequencing data"

### Supplementary Material

Supplementary Figures

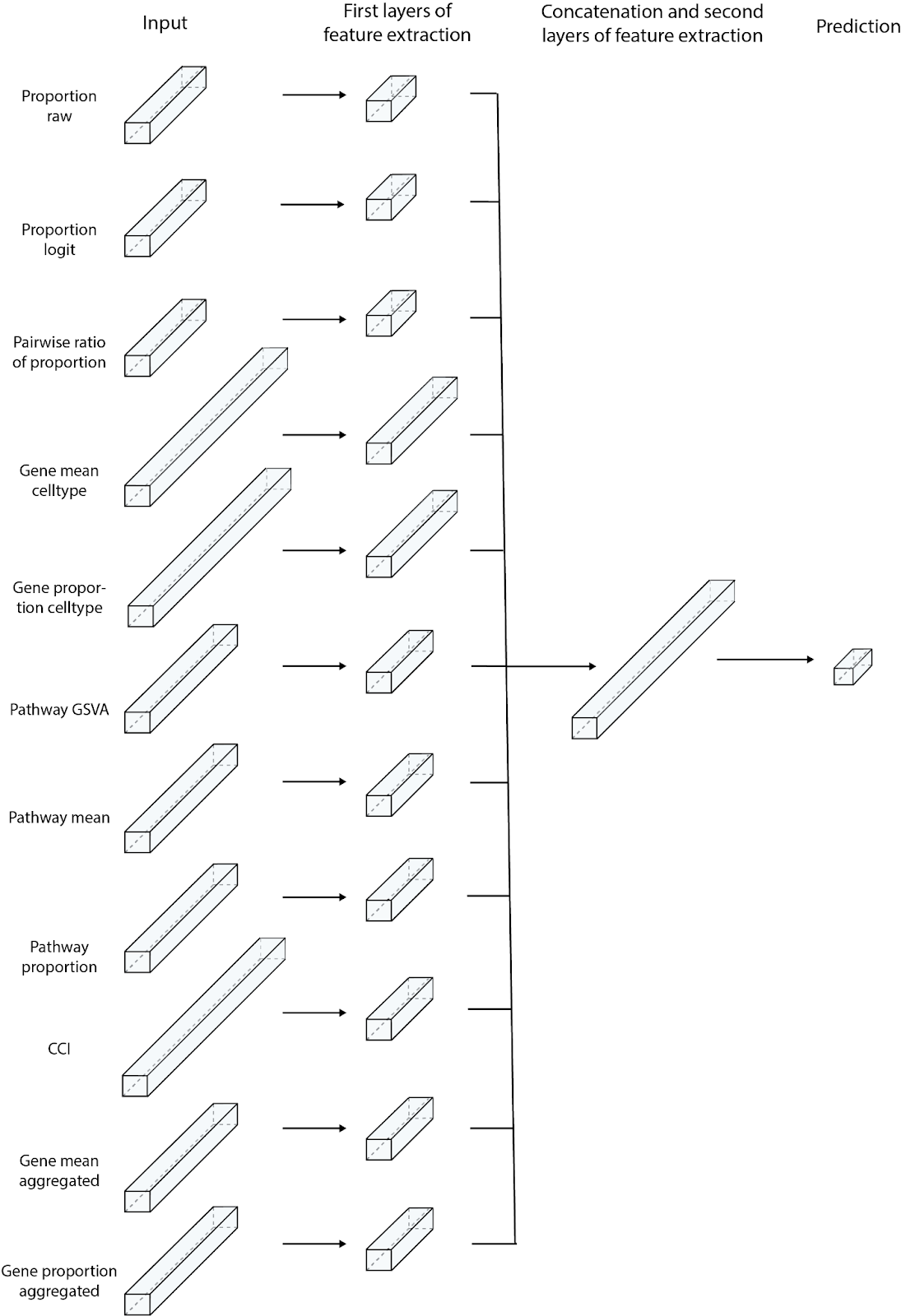

**Supplementary Figure 1. Schematic of the stacked ensemble approach for deep learning.**

The neural network is composed of a total of 11 subnetworks, each taking one feature type as the input and performing feature extraction for that feature type. The extracted features from each feature type are then concatenated and passed through another network for feature extraction and final prediction.

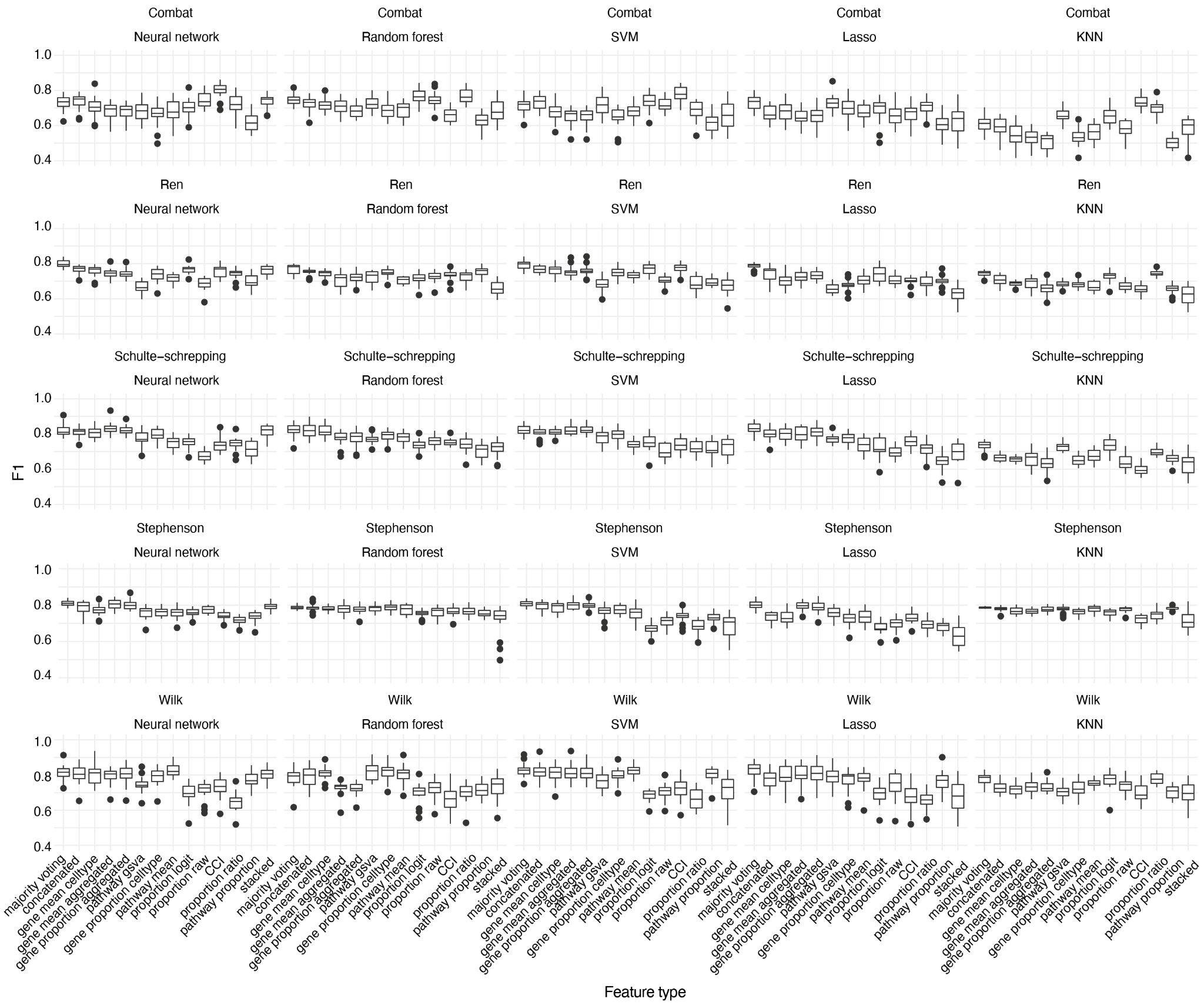

**Supplementary Figure 2. Performance of outcome prediction model on each dataset.**

Each data point in the boxplot represents one F1 score. Each box is made up of 20 F1 scores from the 20 repeated cross-validation performed on the individual dataset.

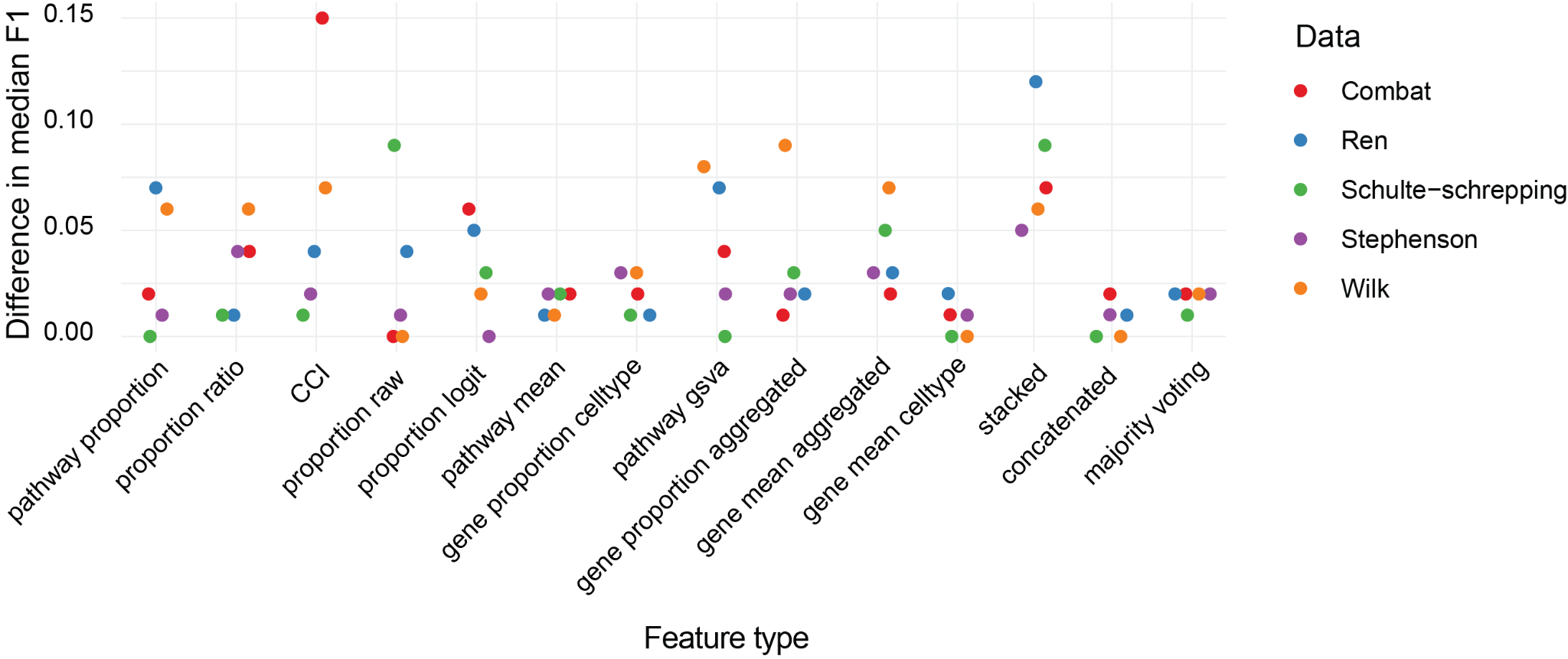

**Supplementary Figure 3. Difference in median F1 between neural network, random forest and SVM.**

For each feature type, we calculate the difference between the maximum and minimum F1 score obtained by neural network, random forest and SVM.

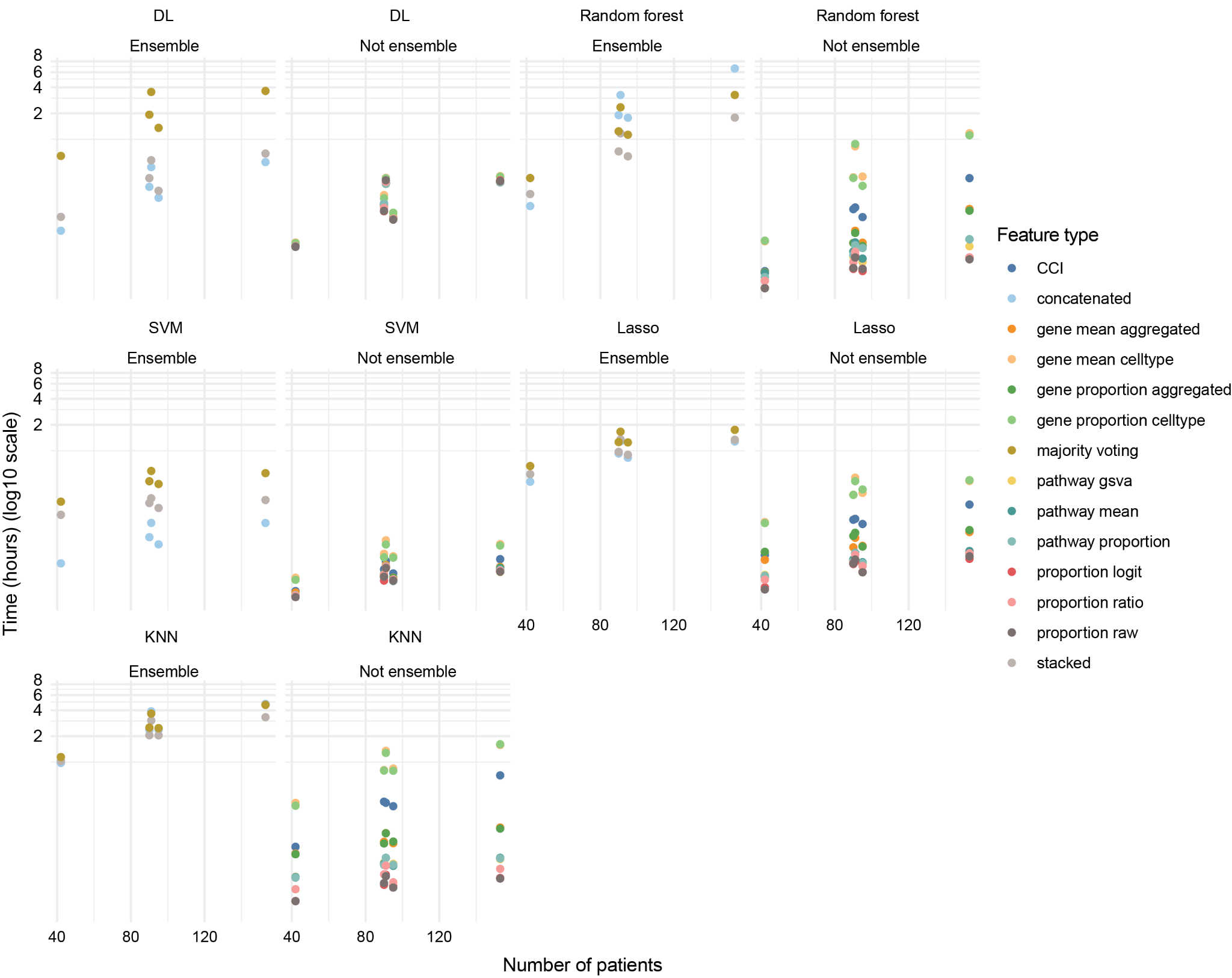

**Supplementary Figure 4. Run time of each feature type for the five COVID-19 datasets.**

Run time was recorded as the CPU time it took to train 20 repeated cross-validation models for each feature type.

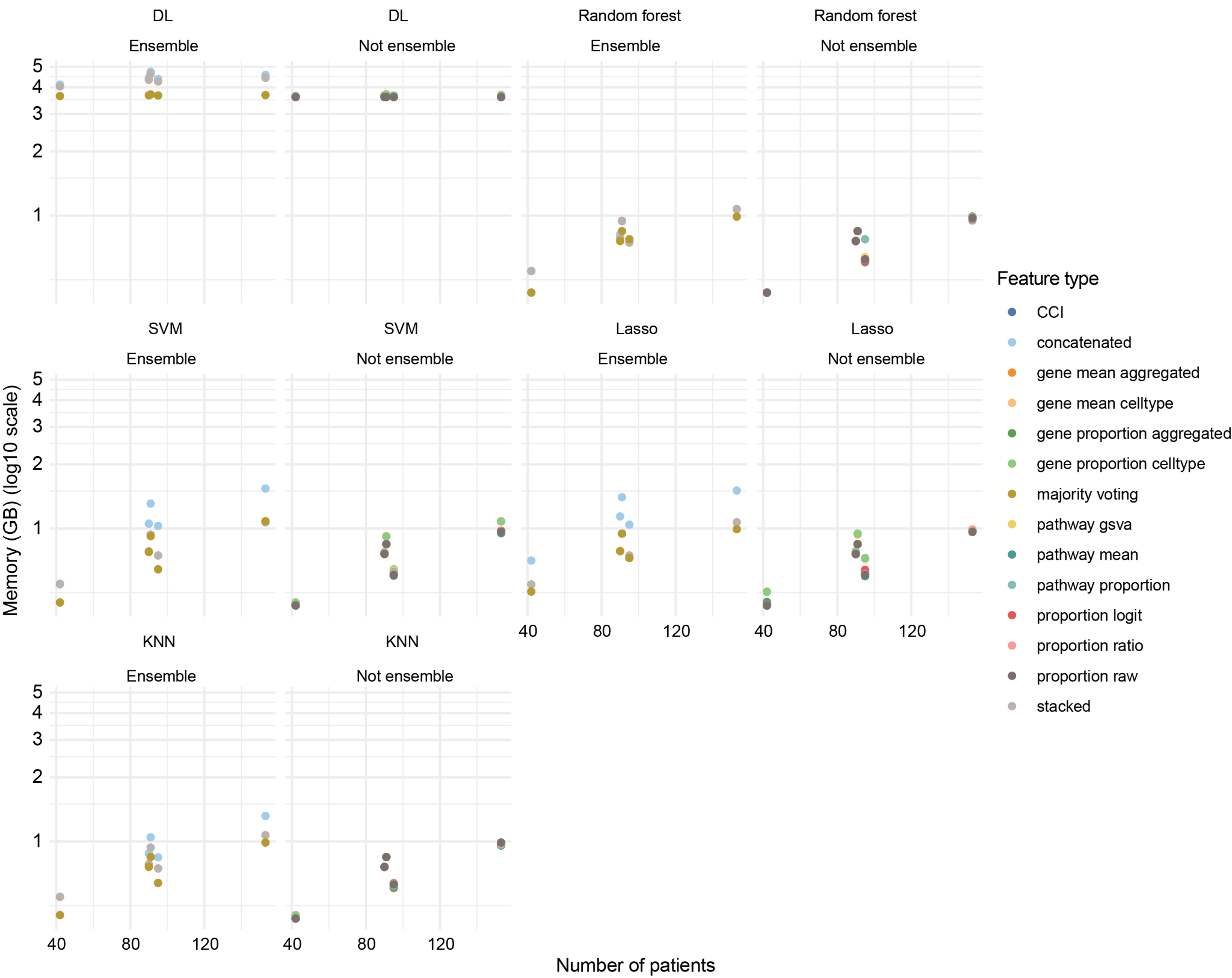

**Supplementary Figure 5. Peak memory usage of each feature type for the five COVID-19 datasets.**

For deep learning model, this was recorded as the sum of peak memory usage from both CPU and GPU. For machine learning models, this was recorded as peak CPU memory.

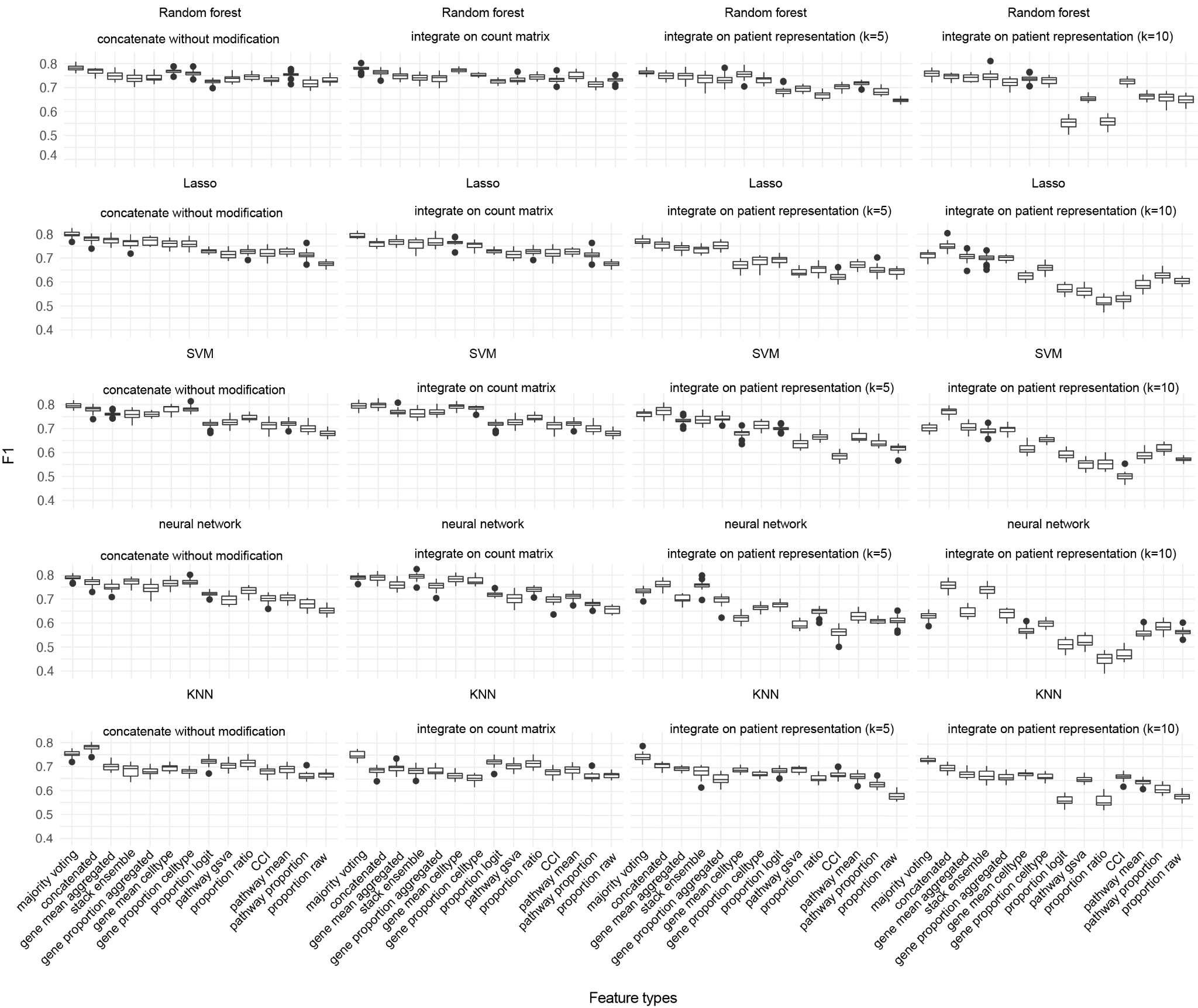

**Supplementary Figure 6. Performance of outcome prediction model on the combination of five datasets.**

Each data point in the boxplot represents one F1 score. Each box is made up of 20 F1 scores from the 20 repeated cross-validation performed on the combined dataset.

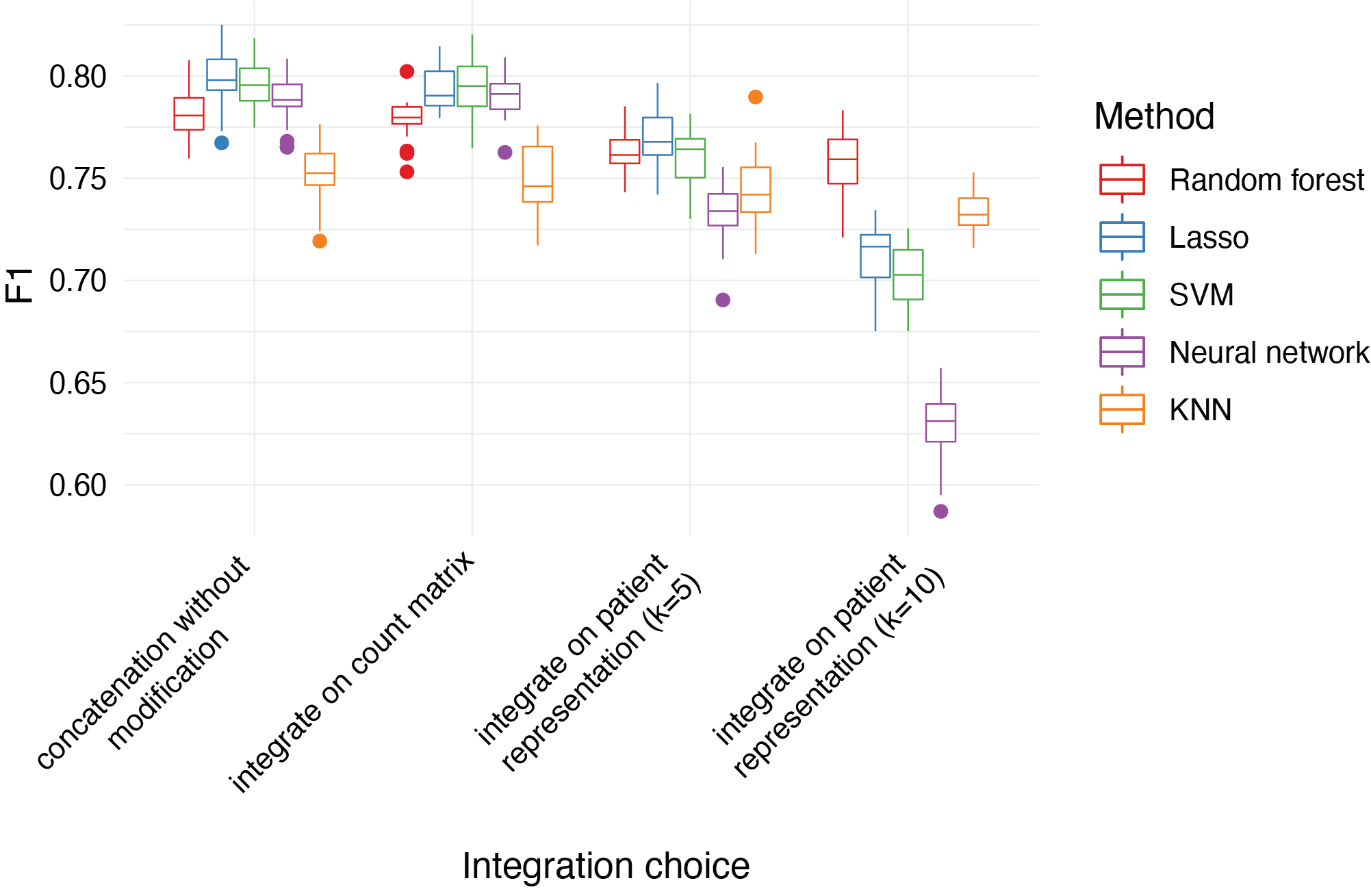

**Supplementary Figure 7. The F1 score for different integration choices and method choices using the ensemble feature type majority voting.**

Each data point in the boxplot represents one F1 score. Each box is made up of 20 F1 scores from repeated cross-validation with 20 repetitions performed on the combined dataset using the ensemble feature type majority voting.

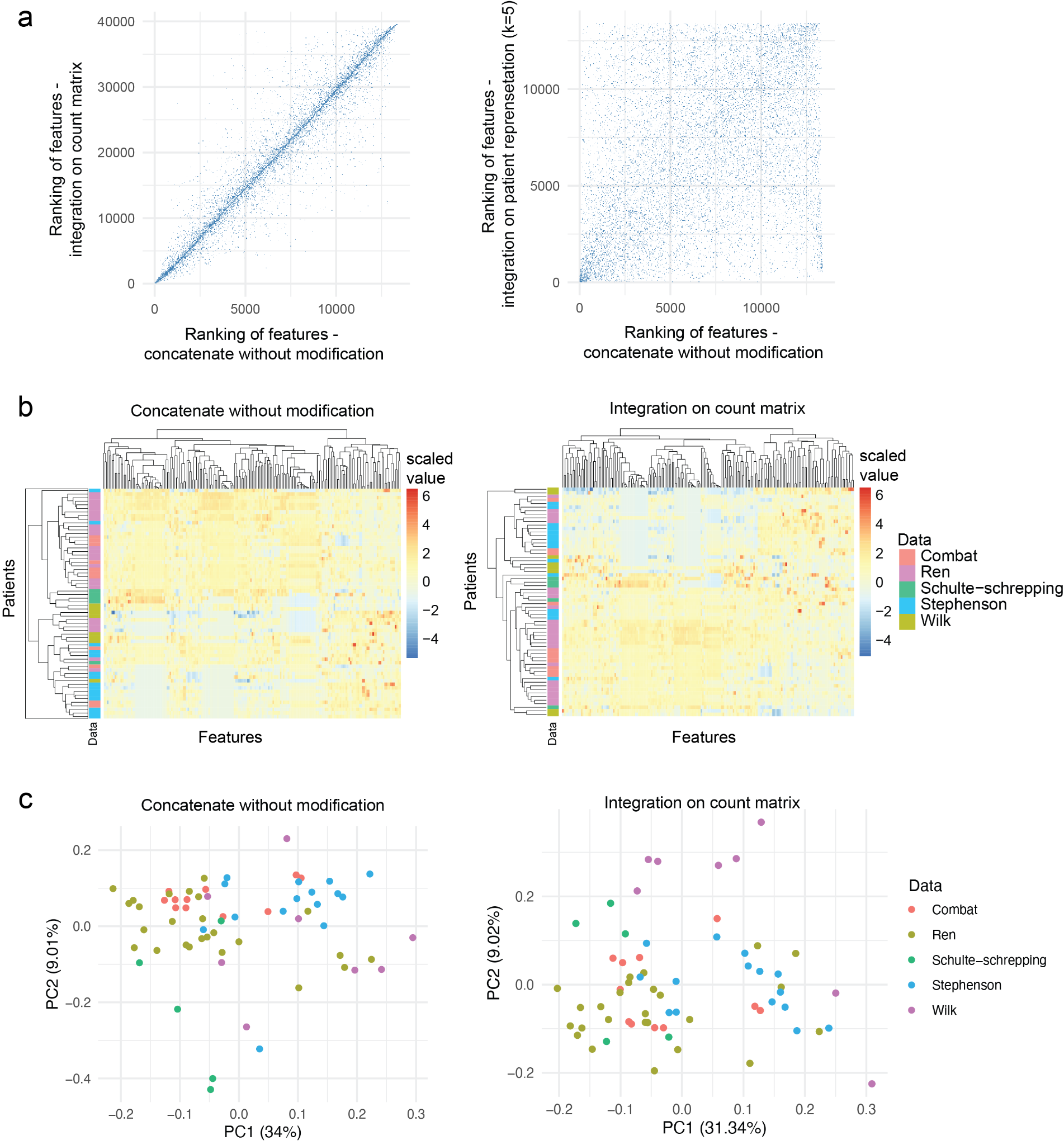

**Supplementary Figure 8. Examination into the features for the different integration choices.** We selected patients in the 41-50 age group, and ran prediction model using the concatenated features as input and SVM as the classification method. We then obtained the rankings of the features based on feature importance score. **a** compares the ranks of the features from different integration choices. The heatmap in **b** plots the values of the top 200 features in each patient. The patient is colour coded by the dataset to reveal any potential batch effect in the features. **c** shows the PCA of the top 200 features, where each dot represents a patient, coloured by the dataset.
